## Supplementary Information for "Heterogeneous recruitment abilities to RNA polymerases generate nonlinear scaling of gene expression with cell volume"

Qirun Wang[1] and Jie Lin[1,2]
[1] *Center for Quantitative Biology, Peking University, Beijing, China*
[2] *Peking-Tsinghua Center for Life Sciences, Peking University, Beijing, China*

### A. Derivation of the binding probability of a promoter

In this section we derive the probability of a promoter bound by an RNAP and we assume that free RNAPs can only bind to unbound promoters with a rate $k_{\text{on}}$. Once the RNAP binds to the promoter, it becomes an initiating RNAP which either unbinds from the promoter with a rate $k_{\text{off}}$ or starts transcribing with a rate $\Gamma_n$. We assume that a transcribing RNAP moves along the gene deterministically and after a gene-dependent time, it drops off the gene and becomes free again. The process is summarized in Figure 1. We assume that the binding probability of promoter $P_b$ is in the steady state and time-independent so that

$$c_n F_n (1 - P_b) k_{\text{on}} = P_b (k_{\text{off}} + \Gamma_n). \tag{S1}$$

Here $c_n$ is the total concentration of RNAPs in the nucleus and $F_n$ is the fraction of free RNAPs. Solving the above equation, we obtain the binding probability as

$$P_{n,b} = \frac{c_n F_n}{c_n F_n + \frac{k_{\text{off}} + \Gamma_n}{k_{\text{on}}}}. \tag{S2}$$

Therefore, the Michaelis-Menten constant is $K_n = \frac{k_{\text{off}} + \Gamma_n}{k_{\text{on}}}$, which is inversely proportional to the binding rate of RNAP to the promoter. A similar model also applies to the translation process and one just needs to replace RNAPs by ribosomes, genes by mRNAs and promoters by ribosome binding sites [1].

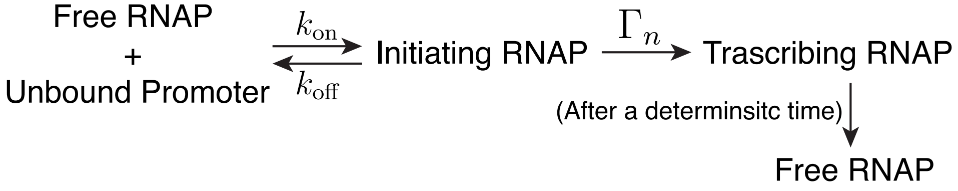


**Supplementary Figure 1** A summary of the transcription process.

### B. Transitions between different phases

In this section, we consider the simple scenario that all genes have the same recruitment ability $1/K_n$ to RNAPs. At the translational level, the protein production rate of gene $i$ is

$$k_{r,i} = \Gamma_r m_i \frac{c_{r,free}}{c_{r,free} + K_r} \tag{S3}$$

where $\Gamma_r$ is the initiation rate of translation, $c_{r,free}$ is the concentration of free ribosomes in the cytoplasm and $K_r$ is the corresponding Michaelis-Menten constant. For simplicity, we assume the recruitment abilities of mRNAs to ribosomes and their translation initiation rates as the same for different genes, therefore the ratios between the protein production rates of different genes are simply equal to the ratios between their mRNA copy numbers. This agrees with the fact that protein levels are often primarily determined by mRNA levels [2, 3] (we will relax this assumption later). Using a similar argument as the transcription process, the conservation of ribosome number leads to

$$\sum_i m_i (1 + \Lambda_{r,i}) \frac{c_r F_r}{c_r F_r + K_r} = r - r F_r. \tag{S4}$$

Here $r$ is the number of ribosomes, $F_r$ is the fraction of free ribosomes. $\Lambda_{r,i} = \Gamma_r L_i / v_r$ is the capacity of ribosomes for mRNA $i$: the maximum number of translating ribosomes a single mRNA can have in the limit when its ribosome binding site is constantly occupied. Here $L_i$ is the length of the mRNA in the unit of codons and $v_r$ is the elongation speed of ribosome.

Therefore, given the translation rate Eq. (S3), we can obtain the time-dependence of non-degradable protein mass

$$\frac{dM_{p,i}}{dt} = k_{r,i}L_i = \Gamma_r m_i \frac{c_{r,free}}{c_{r,free} + K_r} L_i \tag{S5}$$

As the cell mass increases and the cell divides, the ratio between the masses of two proteins will be well approximated by the ratio of their mass production rates. Assuming that the total protein mass $M$ is dominated by non-degradable proteins, we find the mass fraction of protein $i$ in the total proteome in the steady state as

$$\phi_i = \frac{M_{p,i}}{M} = \frac{m_i L_i}{\sum_j m_j L_j}. \tag{S6}$$

Since the lifetimes of mRNAs are short compared with the cell cycle duration, the mRNA numbers are approximately proportional to their production rates, therefore $m_i \propto \Gamma_{n,i} g_i \tau_{m,i}$ and we obtain

$$\phi_i = \frac{M_{p,i}}{M} = \frac{\Gamma_{n,i} g_i \tau_{m,i} L_i}{\sum_j \Gamma_{n,j} g_j \tau_{m,j} L_j}. \tag{S7}$$

Here $\tau_{m,i}$ is the lifetime of the mRNA of gene $i$. Therefore, once the gene copy number is fixed, the mass fraction of one particular protein in the entire proteome is also fixed. In fact, the constant mass fraction is still valid when we relax the assumption that all mRNAs share the same $K_r$ and $\Gamma_r$ and in this case, Eq. (S4) is modified as

$$\sum_i m_i (1 + \Lambda_{r,i}) \frac{c_r F_r}{c_r F_r + K_{r,i}} = r - rF_r, \tag{S8}$$

which can be written as

$$\sum_i \frac{m_i}{V} (1 + \Lambda_{r,i}) \frac{c_r F_r}{c_r F_r + K_{r,i}} = c_r - c_r F_r. \tag{S9}$$

In Phase 1 (see the discussions in the main text and below), $m_i \propto n$ and we assume that the constant mass fractions of proteins are valid so that $n \propto V$, $r \propto V$, which we will confirm self-consistently later. Here $V$ is the cell volume and we assume a constant ratio between the total protein mass and cell volume $\rho = M/V$. Therefore, the above equation is independent of the cell volume and $F_r$ is approximately constant. The ratio between the masses of two proteins becomes

$$\frac{M_{p,i}}{M_{p,j}} = \frac{\Gamma_{r,i} m_i L_i}{\Gamma_{r,j} m_j L_j} \frac{c_r F_r + K_{r,j}}{c_r F_r + K_{r,i}}. \tag{S10}$$

Since $c_r$ is constant as we assume and $F_r$ is constant as we show in the above, we find that Eq. (S10) is also constant once the gene copy number is fixed which self-consistently validates the assumption that the RNAP copy number and ribosome copy number are proportional to the cell volume.

In the following we define the average capacities of genes and mRNAs as

$$\Lambda_n = \sum g_i \Lambda_{n,i} / \sum g_i, \tag{S11}$$

$$\Lambda_r = \sum m_i \Lambda_{r,i} / \sum m_i. \tag{S12}$$

We can rewrite Eq. (S4) as a self-consistent equation to find the mass fraction of free ribosomes in the proteome given the total number of mRNAs and the mass fraction of total ribosomes

$$m_R \frac{\sum_i m_i}{M} (1 + \Lambda_r) \frac{\phi_{r,f}}{\phi_{r,f} + \widetilde{K}_r} = \phi_r - \phi_{r,f}, \tag{S13}$$

here $m_R$ is the protein mass of a single ribosome and $\widetilde{K}_r = K_r m_R / \rho$. Meanwhile, the time derivative of total protein mass is

$$\frac{dM}{dt} = v_r \Lambda_r \frac{\phi_{r,f}}{\phi_{r,f} + \widetilde{K}_r} \sum_i m_i, \tag{S14}$$

and the growth rate is set by the fraction of actively translating ribosomes

$$\mu = \frac{dM/dt}{M} = \frac{v_r}{m_R}\frac{\Lambda_r}{\Lambda_r+1}(\phi_r - \phi_{r,f}). \tag{S15}$$

From Eqs. (S13, S15), we find that to have a constant exponential growth rate as the cell volume increases, the ratio between total mRNA number and total protein mass must be constant. This is satisfied when the transcription rate is proportional to the RNAP number, which is valid when $F_n \ll 1$ and is called Phase 1 of gene expression [4]. From Eq. (8) in the main text, we find that the approximation $F_n \ll 1$ starts to break down when

$$n > \sum_i g_i(1+\Lambda_n) \equiv n_c. \tag{S16}$$

When $n$ is above $n_c$, the linear scaling between the transcription rate and RNAP number begins to break down. In fact, based on the assumption that $c_n \gg K_n$, the transcription rate will quickly be constrained by the gene copy number when Eq. (S16) is satisfied. When the cell enters Phase 2, the growth of cell mass deviates from strict exponential growth. In the extreme scenario, when $c_r F_r \gg K_r$, the translation rate per mRNA will be independent of the ribosome number at all. Assuming $c_r \gg K_r$, one can estimate the transition criteria as

$$r > \sum_i m_i(1+\Lambda_r) \equiv r_c. \tag{S17}$$

### C. Effects of non-specific binding of RNA polymerases

In the main text, we mainly discuss eukaryotic cells in which non-specific binding of RNAPs is not relevant. However, non-specific binding of RNAPs is believed to be significant in bacteria and a finite fraction of DNA-bound RNAPs are found to be inactively transcribing [5]. In this section, we discuss the transition from Phase 1 to Phase 2 taking account of the non-specific binding of RNAPs. In this case, the conservation of total RNAP number becomes

$$\sum_i g_i(1+\Lambda_{n,i})\frac{c_n F_n}{c_n F_n + K_n} + g_{ns}\frac{c_n F_n}{c_n F_n + K_{ns}} = n - nF_n \tag{S18}$$

where $g_{ns}$ is the number of non-specific binding sites and $K_{ns}$ is the non-specific binding Michaelis-Menten constant. In the following, we assume that $K_{ns} \gg K_n$ which we believe is biologically reasonable. One necessary condition for cells to be in Phase 1 is $F_n \ll 1$ therefore we can rewrite Eq. (S18) as

$$\sum_i g_i(1+\Lambda_{n,i})\frac{c_n F_n}{c_n F_n + K_n} + g_{ns}\frac{c_n F_n}{c_n F_n + K_{ns}} = n \tag{S19}$$

To see under what conditions the mRNA production rate is proportional to $n$, we replace $\frac{c_n F_n}{c_n F_n + K_n}$ by $x$ and rewrite the above equation as

$$x + \frac{g_{ns}}{n_c}\frac{K_n}{K_{ns}}\frac{x}{1-x} = \frac{n}{n_c} \tag{S20}$$

where $n_c = \sum_i g_i(1+\Lambda_{n,i})$ and we have used the assumption that $K_{ns} \gg K_n$. We find the condition for $x$ to be proportional to $n$ is that $x \ll 1$. Assuming $g_{ns}K_n/K_{ns}$ is comparable to or smaller than $n_c$, the condition $x \ll 1$ is equivalent to

$$n \ll n_c. \tag{S21}$$

We find that the condition for cells to be in Phase 1 becomes more stringent in the presence of non-specific binding. In the following we focus on Phase 1 and consider a particular gene with the Michaelis-Menten constant $K_{n,i}$ and its transcription rate now becomes

$$k_{n,i} = \Gamma_{n,i} g_i \frac{n}{\frac{K_{n,i}}{K_n}(n_c + g_{ns}\frac{K_n}{K_{ns}}) + (1 - \frac{K_{n,i}}{K_n})n}. \tag{S22}$$

We find that when the cell is in Phase 1 so that $n \ll n_c$, the mRNA production rates of most genes linearly scale with cell volume. The mRNA production rates of those genes with $K_{n,i} < K_n$ can still have a sublinear scaling with cell volume since it is possible to make the two terms in the denominator of Eq. (S22) to be comparable for $n \ll n_c$. However, within Phase 1, the superlinear scaling of those genes with $K_{n,i} > K_n$ is difficult to reach since the first term is always much larger than the second term if $n \ll n_c$.

### D. Protein mass-weighted average of Michaelis-Menten constant

In this section we argue that the appropriate average can be well approximated by the average weighted by the protein mass fractions. Using the self-consistent Eq. (8) in Methods of the main text with heterogeneous $K_{n,i}$ and the definition of $\langle K_{n,i}\rangle$ as the substitution of $K_n$ in Eq. (4) of the main text, we obtain the expression of $\langle K_{n,i}\rangle$ as

$$\frac{\sum_i g_i(1+\Lambda_{n,i})\frac{c_nF_n}{c_nF_n+K_{n,i}}}{\sum_i g_i(1+\Lambda_{n,i})} = \frac{c_nF_n}{c_nF_n+\langle K_{n,i}\rangle}, \tag{S23}$$

therefore,

$$\langle K_{n,i}\rangle = \sum_i \chi_i K_{n,i}, \tag{S24}$$

where

$$\chi_i = \frac{\frac{g_i(1+\Lambda_{n,i})}{c_nF_n+K_{n,i}}}{\sum_j \frac{g_j(1+\Lambda_{n,j})}{c_nF_n+K_{n,j}}}. \tag{S25}$$

We can compare the contributions of two genes as

$$\frac{\chi_i}{\chi_j} = \frac{g_i(1+\Lambda_{n,i})(c_nF_n+K_{n,j})}{g_j(1+\Lambda_{n,j})(c_nF_n+K_{n,i})}. \tag{S26}$$

Meanwhile the ratio of the mRNA levels of two genes is

$$\frac{m_i}{m_j} = \frac{\Gamma_{n,i}g_i\tau_{m,i}(c_nF_n+K_{n,j})}{\Gamma_{n,j}g_j\tau_{m,j}(c_nF_n+K_{n,i})}. \tag{S27}$$

The corresponding ratio of protein mass can be approximated as

$$\frac{M_{p,i}}{M_{p,j}} \approx \frac{\frac{dM_{p,i}}{dt}}{\frac{dM_{p,j}}{dt}} \approx \frac{m_iL_i}{m_jL_j} = \frac{\Lambda_{n,i}g_i\tau_{m,i}(c_nF_n+K_{n,j})}{\Lambda_{n,j}g_j\tau_{m,j}(c_nF_n+K_{n,i})}. \tag{S28}$$

Here we have used $\Lambda_{n,i} = \Gamma_{n,i}L_i/v_n$. Comparing Eqs. (S26) and (S28), we find that these two ratios are highly correlated, validating our choice of the weight as the protein mass fraction.

### E. Heterogeneous initiation rates

We extend our simulations in the main text (Figure 3a, b) to include heterogeneous initiation rates. We first computed the initiation rates that are constant for genes except RNAP and ribosome, using the same protocol as the one introduced in Methods of the main text. We then added noises to the initiation rates $\Gamma_{n,i}$ so that the distribution becomes a lognormal distribution with the the average the same as the constant value before adding noises. The coefficient of variation (CV) of the distribution is equal to 1. We then added the effects of fluctuating initiation rates to the Michaelis-Menten constant $K_{n,i}$ as $K_{n,i} = K^0_{n,i} + \frac{\Gamma_{n,i}}{k_{on}}$ where $k_{on} = 0.01$ and $K^0_{n,i}$ follows a lognormal distribution with mean equal to $6 \times 10^3/\mu m^3$ and CV equal to 0.5.

We also simulated a modified model to mimic a scenario in which the nonlinear scaling has nothing to do with the recruitment abilities. We considered two sets of Michaelis-Menten constants such that they share the same randomness of the heterogeneous initiation rates $K_{n,i} = K^0_{n,i} + \frac{\Gamma_{n,i}}{k_{on}}$, but their random off-rates $k_{off,i}$ are independent of each other so that their $K^0_{n,i}$ are different. The simulation is the same as Figure 6b, except that the nonlinear degrees and the mRNA production rates are calculated using the two sets of $K_{n,i}$ respectively to simulate the case that the nonlinear scaling is independent of the Michaelis-Menten constants. We found that the Pearson correlation coefficient between the nonlinear scalings and mRNA production rates becomes negative rather than positive (Figure 6d), in contrast to the original model. We repeated the simulation multiple times and find that the correlation coefficient in the modified model is always smaller than that of the original model (Figure 6e).

### F. Simulations of periodic cell cycle

In this section, we simulated the case of periodic cell cycle. We chose two degradable proteins, one superlinear and one sublinear, and used the ratio of their concentrations to determine the timing of cell division. When the ratio of their concentrations exceeds some threshold value, the cell divides, in concert with the idea that the ratio of cell cycle regulators determines cell division [6]. Note that we also have an additional requirement on the minimum cell-cycle duration to avoid cell division immediately after cell birth.

We considered symmetric division so that all mRNAs and proteins are symmetrically distributed between the two daughter cells. For simplicity, we assumed that the Michaelis-Menten constants of RNAP binding and ribosome binding are constant in time and constant gene copy numbers. Note that our qualitative results are independent of these assumptions and in more realistic models, these time dependences must be considered. We tracked a single lineage of cell so that we monitored one of the daughter cells after cell division. In the simulations, we took the lifetimes of mRNAs and degradable proteins as 5 mins. The sublinear and superlinear proteins that we used as the signal proteins respectively have $K_{n,i} \approx 970$ and $K_{n,i} = 2.8 \times 10^4$. The cell divides when the ratio of the superlinear protein concentration and the sublinear protein concentration exceeds 0.3. Other simulation details are the same as Figure 3a, b in the main text.

We found that for superlinear genes, their mRNA and protein concentrations decrease initially at the beginning of the cell cycle due to the halved RNAP number at cell birth, but quickly increases as the RNAP number increases (vice versa for sublinear genes). As the cell gets the periodic steady state, all mRNAs and proteins double their numbers at cell division compared with cell birth (Figure 12).

Table S1: A summary of the parameters used in the numerical simulations if not mentioned in the text. Note that some of the parameters may not be realistic estimations of any specific organisms and our main conclusions are independent of the chosen parameters.

| Parameters | Meaning | Values |
|---|---|---|
| $\mu_0$ | attempted growth rate | 0.5 division per hour |
| $\tau_{p,i}$ | lifetime of degradable proteins | 10 min |
| $v_r$ | translational speed of ribosome | 12 aa/sec |
| $v_n$ | transcriptional speed of RNAP | 36 nt/sec |
| $N$ | number of genes | 2000 |
| $L_r$ | number of amino acids of ribosome | $10^4$ |
| $L_n$ | number of amino acids of RNAP | $10^3$ |
| $L_i$ | number of amino acids of other proteins | 500 |
| $\Gamma_r$ | initiation rate of translation | 10 $min^{-1}$ |
| $K_r$ | Michaelis-Menten constant of ribosome binding | $6000/\mu m^3$ |
| $\rho$ | ratio between total protein mass and cell volume | $10^{10}$ $aa/\mu m^3$ |
| $a$ | ratio between cell volume and nuclear volume | 20 |

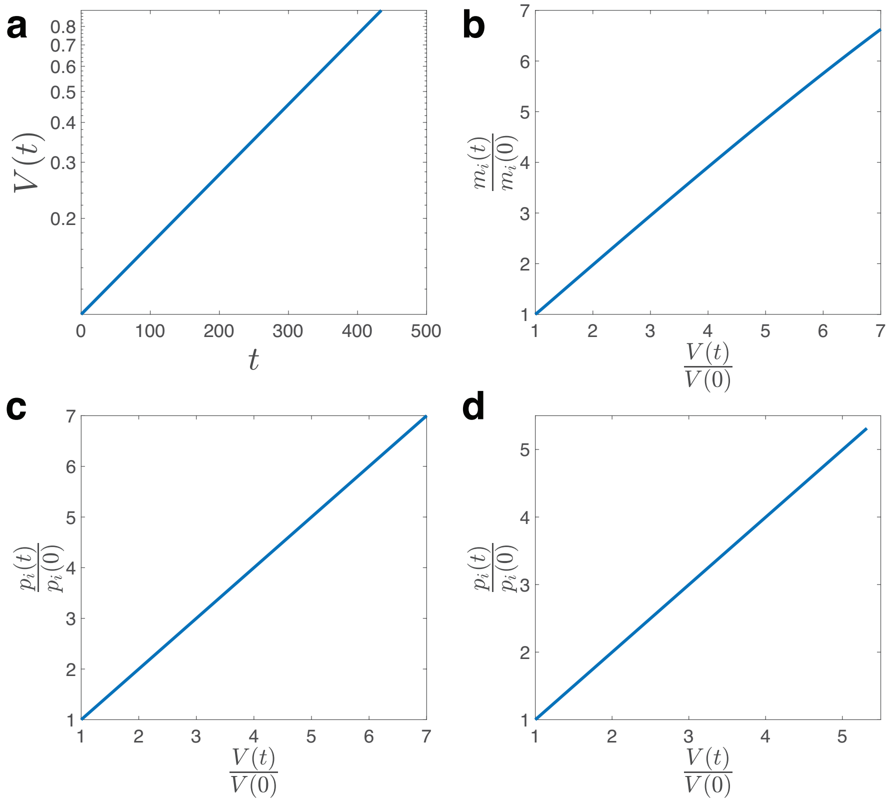


**Supplementary Figure 2** Numerical simulations of the homogeneous model in which all genes share the same recruitment ability to RNAPs. (a) The cell volume grows exponentially, which is proportional to the total protein mass. (b) The mRNA copy number is proportional to the cell volume. (c) The protein number increment is proportional to the cell volume increment for non-degradable proteins. (d) The protein copy number is proportional to the cell volume for degradable proteins.

Table S2: Transcription factors binding motifs annotated in Yeastract database[7] enriched in the sublinear regime.

| TF Gene | ORF | Motif |
|---|---|---|
| ABF2 | YMR072W | |
| ACA1 | YER045C | |
| ARG80 | YMR042W | |
| ARR1 | YPR199C | |
| CAD1 | YDR423C | |
| CIN5 | YOR028C | |
| CST6 | YIL036W | |
| ECM22 | YLR228C | |
| FHL1 | YPR104C | |
| FKH1 | YIL131C | |
| FZF1 | YGL254W | |
| GAT3 | YLR013W | |
| GCN4 | YEL009C | |
| GCN4 | YEL009C | |
| GCN4 | YEL009C | |
| GCN4 | YEL009C | |
| GCN4 | YEL009C | |
| GCN4 | YEL009C | |
| GCN4 | YEL009C | |
| GCN4 | YEL009C | |
| GIS1 | YDR096W | |
| GLN3 | YER040W | |
| GLN3 | YER040W | |
| HSF1 | YGL073W | |
| HSF1 | YGL073W | |
| HSF1 | YGL073W | |
| INO2 | YDR123C | |
| INO4 | YOL108C | |
| IXR1 | YKL032C | |
| KAR4 | YCL055W | |
| MATALPHA2 | YCR039C | |
| MCM1 | YMR043W | |
| MCM1 | YMR043W | |
| MGA1 | YGR249W | |
| MGA1 | YGR249W | |
| MOT3 | YMR070W | |
| MOT3 | YMR070W | |
| MOT3 | YMR070W | |
| MOT3 | YMR070W | |

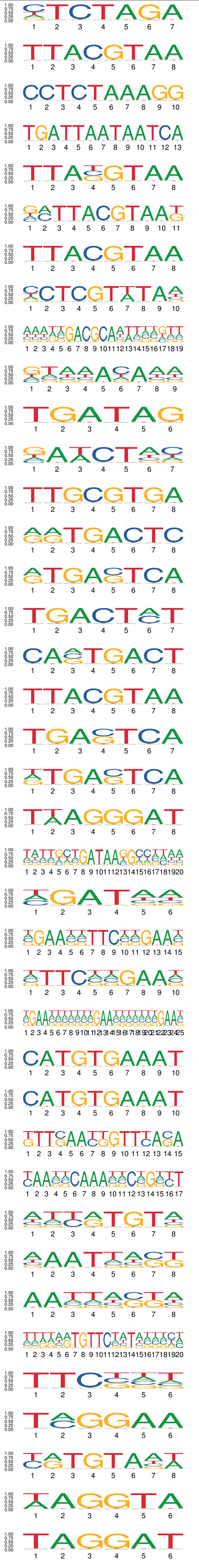

| TF Gene | ORF | Motif |
|---|---|---|
| MOT3 | YMR070W | |
| MOT3 | YMR070W | |
| MSN2 | YMR037C | |
| PDR8 | YLR266C | |
| RLM1 | YPL089C | |
| RME1 | YGR044C | |
| SFL1 | YOR140W | |
| SIP4 | YJL089W | |
| SKO1 | YNL167C | |
| SKO1 | YNL167C | |
| SMP1 | YBR182C | |
| SRD1 | YCR018C | |
| SRD1 | YCR018C | |
| STB5 | YHR178W | |
| STE12 | YHR084W | |
| SUM1 | YDR310C | |
| TBF1 | YPL128C | |
| TEC1 | YBR083W | |
| TEC1 | YBR083W | |
| UPC2 | YDR213W | |
| URC2 | YDR520C | |
| VHR1 | YIL056W | |
| XBP1 | YIL101C | |
| XBP1 | YIL101C | |
| XBP1 | YIL101C | |
| YAP1 | YML007W | |
| YAP1 | YML007W | |
| YAP1 | YML007W | |
| YAP1 | YML007W | |
| YAP3 | YHL009C | |
| YAP3 | YHL009C | |
| YAP3 | YHL009C | |
| YAP3 | YHL009C | |
| YAP6 | YDR259C | |
| VHR2 | YER064C | |
| COM2 | YER130C | |
| YOX1 | YML027W | |
| YPR015C | YPR015C | |

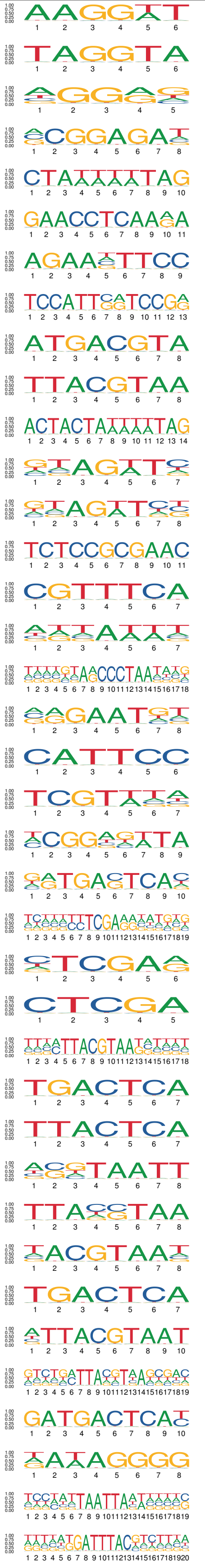

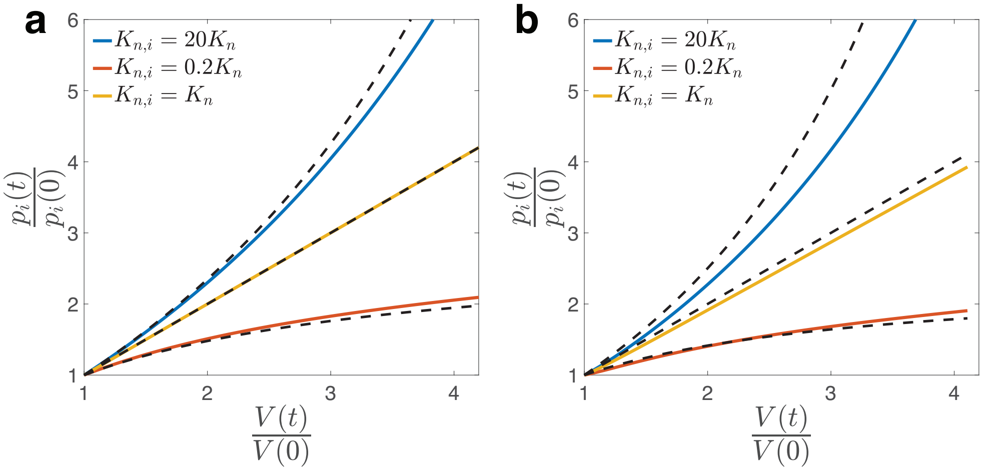


**Supplementary Figure 3** Numerical simulations of the simple model in which all genes share the same recruitment ability to RNAPs except two genes. We show the scaling behaviors of degradable proteins here with lifetime $\tau_{p,i} = 10$ min (a) and $\tau_{p,i} = 40$ min (b). The dashed lines are the theoretical predictions which are the same as those of mRNAs. In (a), we take the simulation results at $t = 100$ as the initial values to remove the transient effects and. In (b), we take the simulation results at $t = 150$ as the initial values.

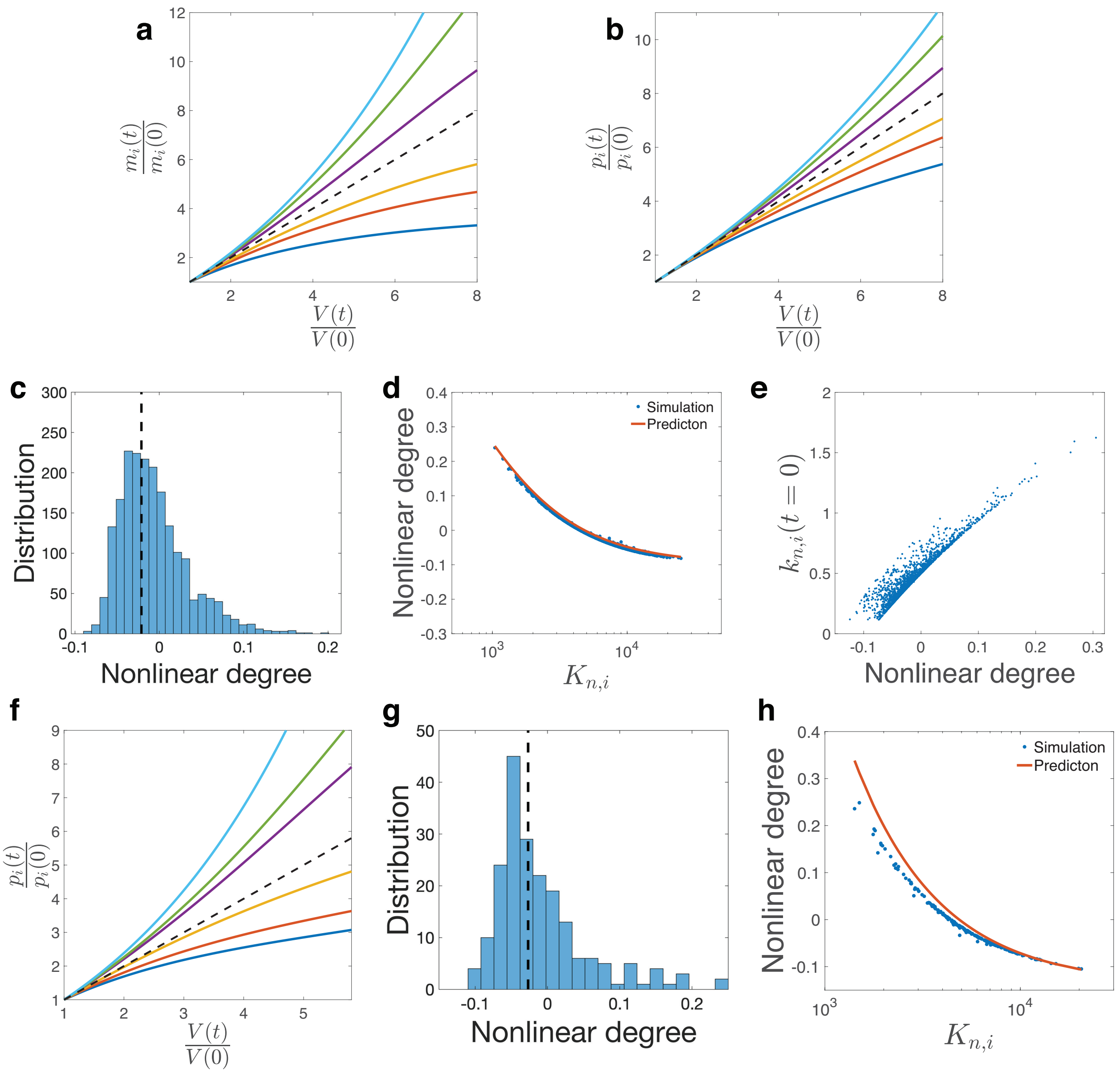


**Supplementary Figure 4** Numerical simulations of the full model in which $K_{n,i}$ is continuously distributed. (a) Examples of mRNA numbers *vs.* cell volume including superlinear and sublinear scaling. The dashed line is the $y = x$ line. (b) Examples of non-degradable protein number *vs.* cell volume. The dashed line is the $y = x$ line. (c) Distribution of the measured nonlinear degrees $\alpha$ of non-degradable protein numbers from numerical simulations. The dashed line marks the location of the median value of the nonlinear degrees. (d) We compare the theoretically predicted nonlinear degrees of protein numbers and the measured values from numerical simulations. (e) The mRNA production rate at time zero *v.s.* their corresponding nonlinear degrees. (f) We show some examples of the scaling behaviors of degradable proteins. The dashed line is the $y = x$ line. (g) Distribution of the measured nonlinear degrees $\beta$ of degradable protein numbers from numerical simulations. The dashed line marks the location of the median value of the nonlinear degrees. (h) We compare the theoretically predicted nonlinear degrees of degradable proteins and the measured values from numerical simulations.

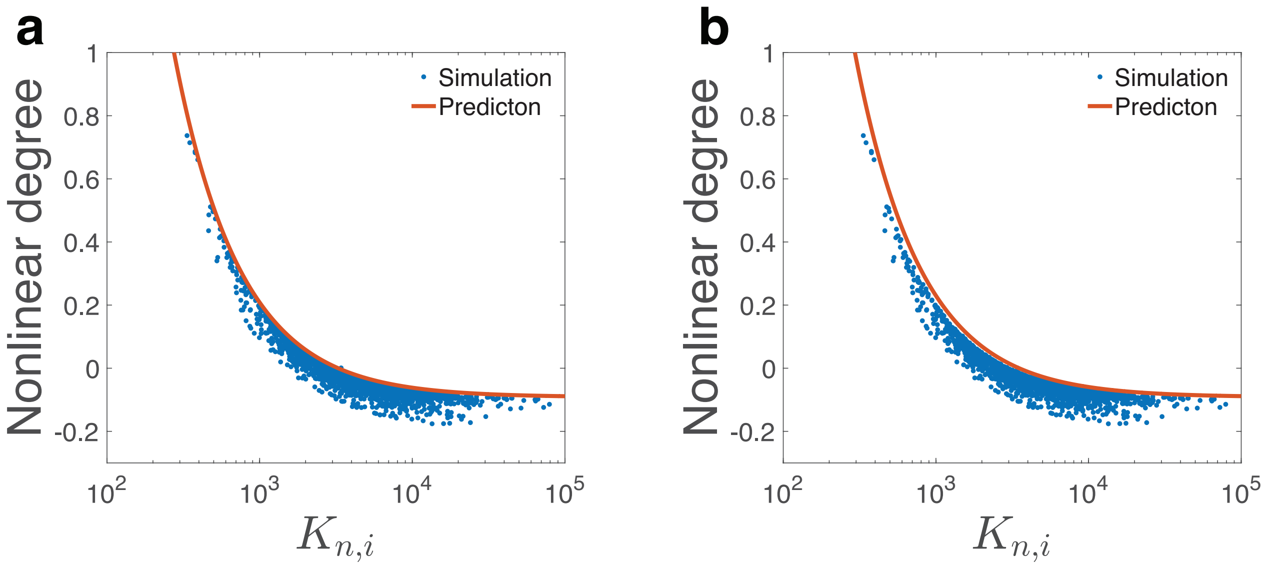


**Supplementary Figure 5** We compare the theoretically predicted nonlinear degrees of mRNA numbers and the measured one from numerical simulations. In both panels, the coefficient of variation of the MM constants is 1. (a) The average MM constant is computed as a weighted average over the initial protein mass fractions. (b) The same simulations as the left panel, but the weight is based on the time-averaged protein mass fractions over the total duration of simulation.

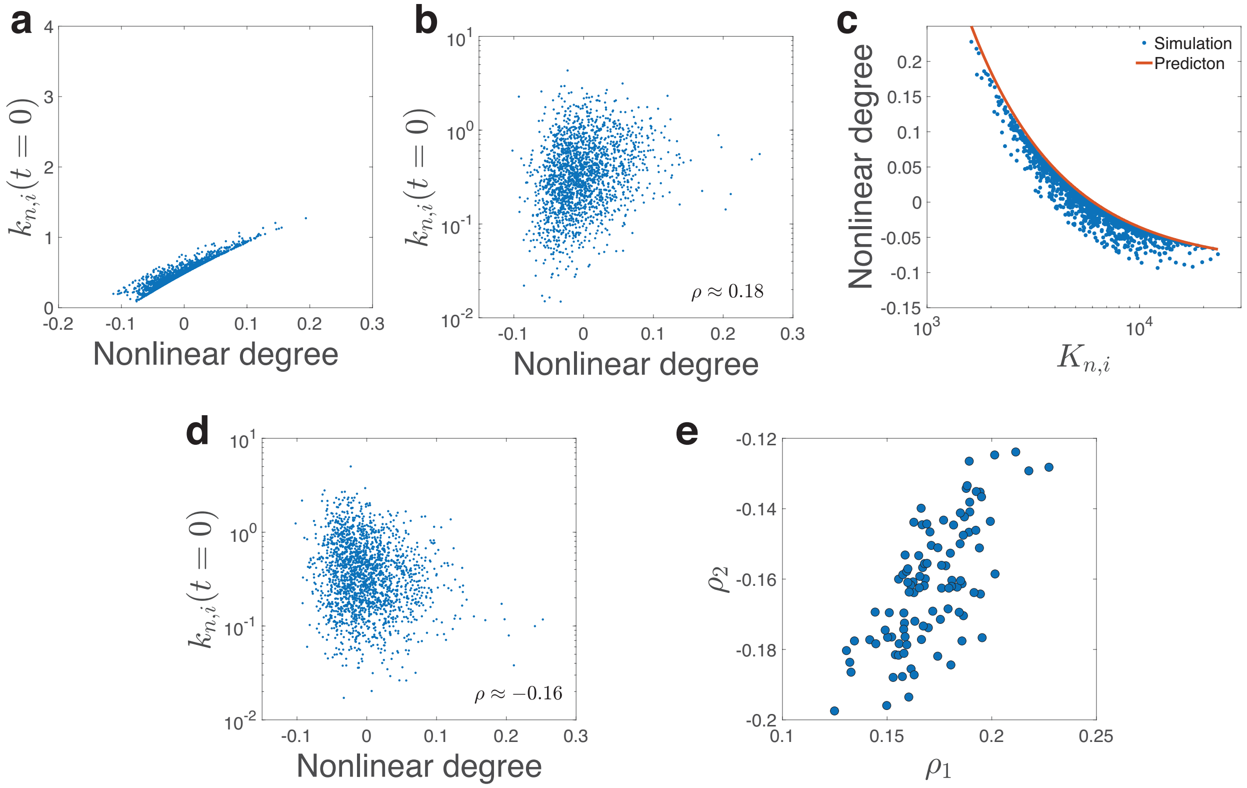


**Supplementary Figure 6** We simulate the more general model in which $K_{n,i} = (k_{off,i} + \Gamma_{n,i})/k_{on}$. The p values are generated from the two-sided Pearson correlation test. (a) In this panel, $\Gamma_{n,i}$ is constant (except for ribosome and RNAP). The Pearson correlation coefficient between the mRNA production rates (y axis) and the nonlinear degrees of mRNA numbers is 0.86 with the p value $<$ 2.20e-16. (b) We add heterogeneity to $\Gamma_{n,i}$ so that its coefficient of variation (standard deviation/mean) is 1. The Pearson correlation coefficient is reduced to 0.18 with the p value = 3.27e-16, close to the experimental value, 0.17 (Figure 4b in the main text). (c) We confirm that our main results on the relation between the nonlinear degree and the Michaelis-Menten constant are still valid in the presence of heterogeneity in $\Gamma_{n,i}$. (d) The negative correlation between the mRNA production rates and the nonlinear scaling for the model in which the recruitment abilities do not affect the nonlinear scaling. The Pearson correlation coefficient is reduced to $-0.16$ with the p value = 4.02e-13. (e) We repeat the simulations multiple times and compare the correlation coefficients of the two models. The correlation coefficients of the model in which the recruitment ability does not influence the nonlinear scaling ($\rho_2$) are negative and always smaller than those of our model ($\rho_1$).

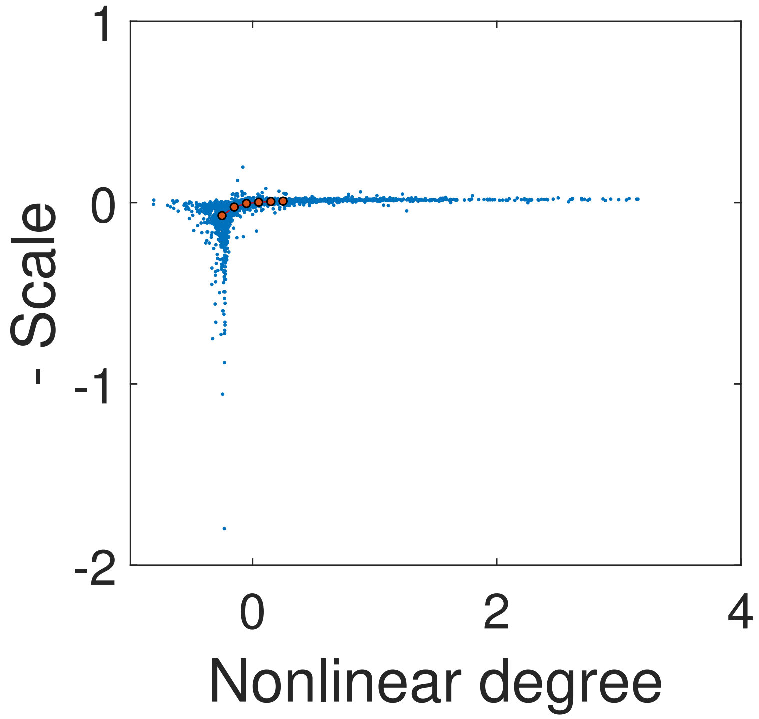


**Supplementary Figure 7** The y axis is the minus nonlinear scale in Ref. [6] calculated as the slope in the linear fitting of mRNA concentration *vs.* cell volume. The x axis is the nonlinear degree $\beta$ defined in this work. The Spearman correlation coefficient is 0.8741426 (two-sided Spearman correlation test, p value $<$ 2.20e-16). The red points are the median values after binning.

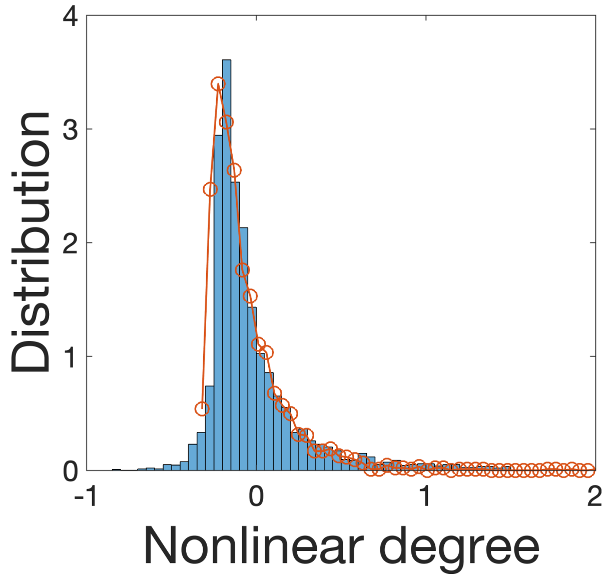


**Supplementary Figure 8** We compare the numerically simulated distribution of nonlinear degrees (circles) and the experimentally measured distribution. In the simulation, the distribution of the Michaelis-Menten constants follow a lognormal distribution with a coefficient of variation equal to 0.8. $n_c = 3200$ and the simulations stop when the total protein mass are larger than $2.5M_b$. Other simulation details are the same as Figure 3 in the main text.

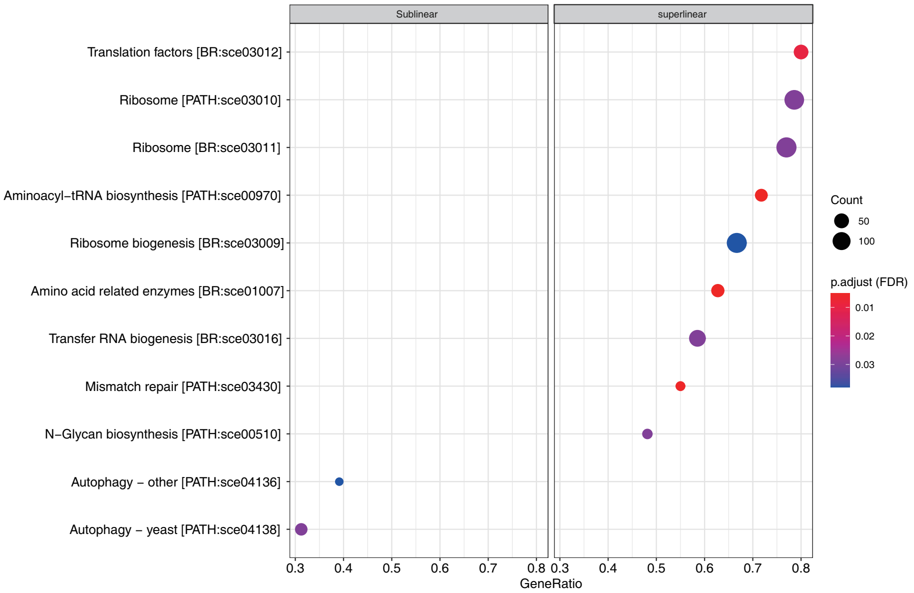


**Supplementary Figure 9** Functional gene sets enriched in the nonlinear scaling regime. GeneRatio represents tags in GSEA, which is the fraction of leading-edge genes in genes both occurring in our list and in the corresponding gene set. Point size represents the number of leading-edge genes. Single-sided permutation test of GSEA was performed and colors of the points represent the adjusted p value (FDR). Names of the gene sets are followed by their IDs in KEGG data base[8–10].

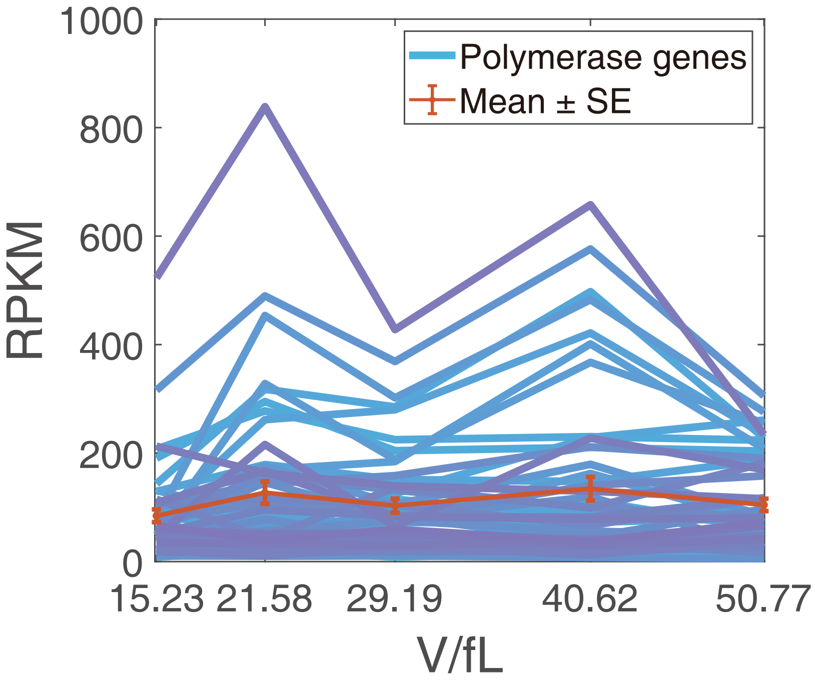


**Supplementary Figure 10** The RPKM values of RNA polymerase II genes and their average value. There are 52 genes annotated as RNA polymerase II holoenzyme in Gene Ontology database using AmiGO [11–13]. Error bars represent standard errors (SE). Two-sided Wilcoxon test results show no significant between-groups differences (V1 *vs.* V2: W = 1196, p value = 3.12e-1; V2 *vs.* V3: W = 1390, p value = 8.07e-1; V3 *vs.* V4: W = 1357, p value = 9.77e-1; V4 *vs.* V5: W = 1312, p value = 7.97e-1).

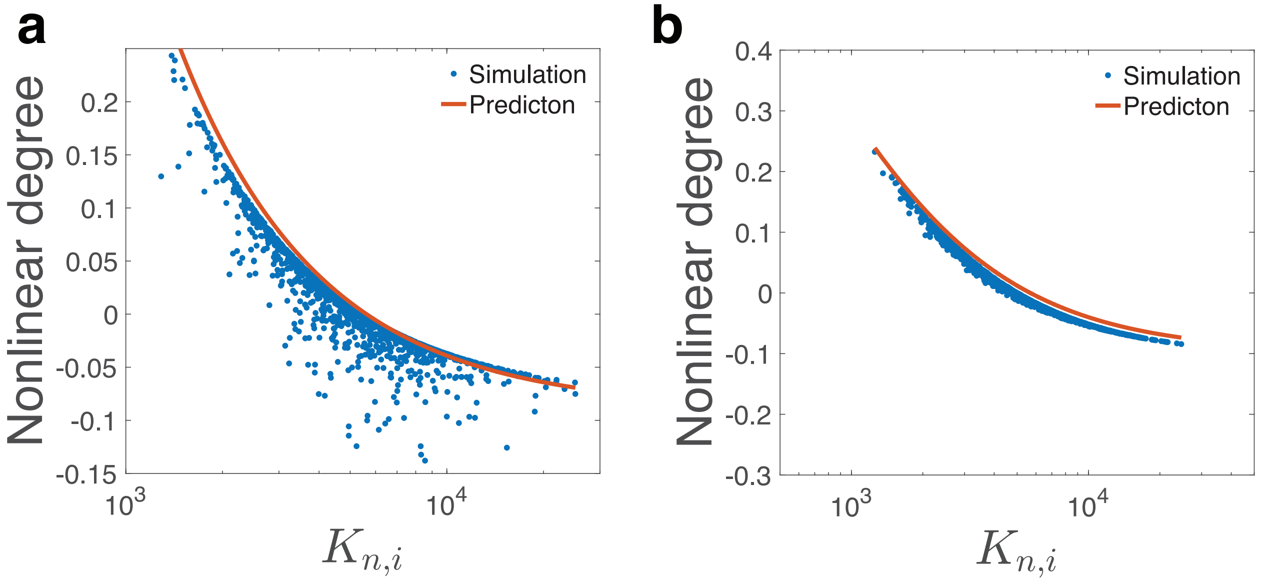


**Supplementary Figure 11** Numerical simulations of the full model in which $K_{n,r}$ is larger than $\langle K_{n,i} \rangle$. The nonlinear degree $\beta$ of the ribosome gene is about $-0.2$. (a) We compare the theoretically predicted nonlinear degrees of mRNA numbers and the measured values from numerical simulations. (b) We compare the theoretically predicted nonlinear degrees of protein numbers and the measured values from numerical simulations.

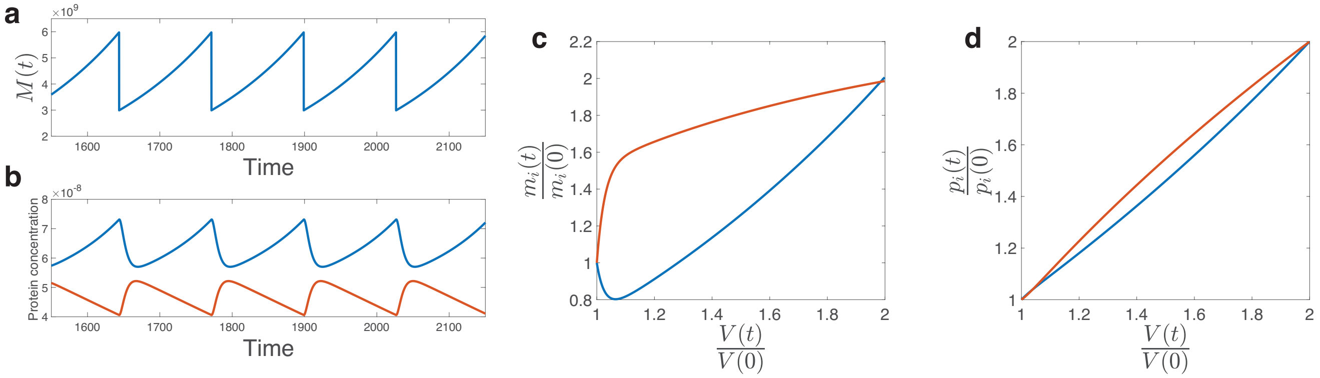


**Supplementary Figure 12** Simulations of periodic cell cycle. (a) The time trajectory of total protein mass in the periodic cell cycle simulation. (b) The time trajectory of the two signal proteins, one superlinear (blue) and one sublinear (red). The cell divides when the ratio of their concentrations exceeds some threshold value. Note that the y axis of the blue curve is multiplied by six to better illustrate the data. (c) The volume-dependence of superlinear (blue) and sublinear (red) mRNA numbers. (d) The same analysis as (c) for non-degradable proteins.

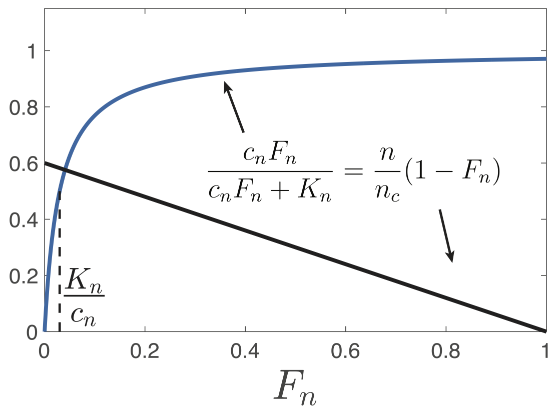


**Supplementary Figure 13** Based on the conservation of the total number of RNAPs, a self-consistent equation, as shown in the figure, is derived for the case of homogeneous recruitment abilities to RNAPs (Eq. (8) in Methods of the main text). Here $n$ is the total number of RNAPs, $n_c = \sum_i g_i(1 + \Lambda_{n,i})$, and $F_n$ is the fraction of free RNAPs. $c_n$ is the concentration of total RNAPs in the nucleus. All variables except $F_n$ are given. The blue curve and the black line are respectively the left and right sides of the equation shown in the figure. The intersection of the two curves allows us to find the $F_n$ that solves the self-consistent equation. Assuming $K_n/c_n \ll 1$, the fraction of free RNAPs $F_n$ that solves Eq. (8) in Methods must be much smaller than 1 if $n < n_c$.
